# The glutathione cycle shapes synaptic glutamate activity

**DOI:** 10.1101/325530

**Authors:** Thomas W. Sedlak, Bindu D. Paul, Greg M. Parker, Lynda D. Hester, Yu Taniguchi, Atsushi Kamiya, Solomon H. Snyder, Akira Sawa

## Abstract

Glutamate is the most abundant excitatory neurotransmitter, present at the bulk of cortical synapses, and participating in many physiologic and pathologic processes ranging from learning and memory to stroke. The tripeptide, glutathione, is one third glutamate and present at up to low millimolar intracellular concentrations in brain, mediating antioxidant defenses and drug detoxification. Because of the substantial amounts of brain glutathione and its rapid turnover under homeostatic control, we hypothesized that glutathione is a relevant reservoir of glutamate, and could influence synaptic excitability. We find that drugs which inhibit generation of glutamate by the glutathione cycle elicit decreases in cytosolic glutamate and decreased miniature excitatory post synaptic potential (mEPSC) frequency. In contrast, pharmacologically decreasing the biosynthesis of glutathione leads to increases in cytosolic glutamate and enhanced mEPSC frequency. The glutathione cycle can compensate for decreased excitatory neurotransmission when the glutamate-glutamine shuttle is inhibited. Glutathione may be a physiologic reservoir of glutamate neurotransmitter.

**Significance:** Glutathione is the principal antioxidant and redox regulator in cells. In addition to its essential roles in redox homeostasis it functions as cofactors for a multitude of enzymes. We show here that glutathione is a reservoir for synaptic glutamate, the excitatory neurotransmitter in the central nervous system. Deficits in glutathione have been linked to multiple neurodegenerative and neuropsychiatric disorders. Accordingly, agents that restore glutathione-glutamate homeostasis may afford therapeutic benefit.

---

Glutamate is the most abundant excitatory transmitter in the central nervous system, utilized at 50- 70% of cortical synapses (1, 2). Glutamate participates in diverse physiological processes, such as developmental plasticity and long-term potentiation as well as brain diseases: epilepsy, stroke, amyotrophic lateral sclerosis, Alzheimer’s disease, Parkinson’s disease and schizophrenia (3). Glutathione is a tripeptide of glutamate, cysteine and glycine, occurring in neurons at concentrations of 0.2-2 mM; it is the most abundant low molecular weight thiol of bacteria, plant and animal cells (4-6). As such, it regulates critical cellular processes such as metabolism of estrogens, prostaglandins, leukotrienes and xenobiotic drugs. Glutathione is well known as an anti-oxidant agent, providing a major line of defense against oxidative and other forms of stress, largely as a cofactor for the glutathione peroxidase and S-transferase enzyme families (7-10).

Glutathione metabolism is governed by the glutathione cycle (Fig. 1*A*, Fig. S1), in which glutamate is added and liberated at discrete steps (4, 11). Glia serve as a major supplier of cysteine for neuronal glutathione synthesis, and 50-60% of glutamate neurotransmitter is derived from the glutamine-glutamate shuttle between neurons and glia, with substantially smaller amounts of glutamate transmitter derived from glycolysis (12-14). However, this shuttle is not the only means to replenish supply of neuronal glutamate; when it is inhibited, neurons quickly restore glutamate neurotransmission by an ill-defined endogenous mechanism, suggesting that neurons might be making use of a storage buffer of glutamate (15). We hypothesize that the glutathione cycle may be one such glutamate reserve, especially considering its high concentration and short half-life.

**Fig. 1.**
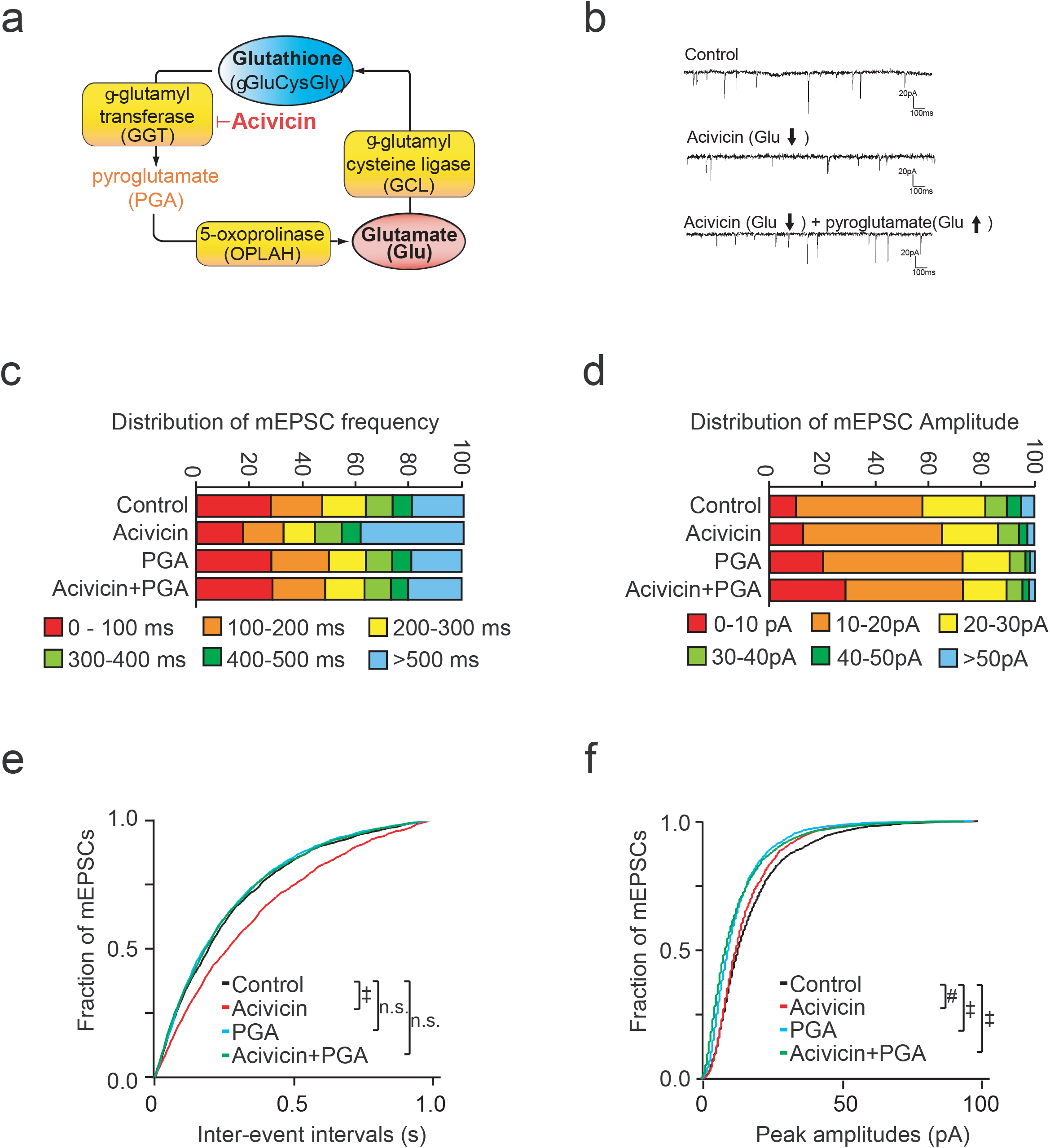
Blocking efflux of glutamate from the glutathione cycle decreases mEPSC. (*A*) Schematic representation of the glutathione cycle and inhibition of gamma-glutamyl transferase (GGT) by acivicin, which is upstream of the liberation of glutamate. Details appear in Supplementary Figures 1 and 2. (*B*) Representative mEPSC traces in primary cortical neurons treated 24 h with vehicle, 25 μM acivicin and/or 5 μM pyroglutamate (PGA). Acivicin decreased mEPSC frequency which can be recovered by pyroglutamate. (*C*) Distribution of mEPSC frequency in cortical neurons treated with acivicin. (*D*) Distribution of mEPSC amplitude in cortical neurons treated with acivicin. (*E*) Cumulative probability plots of mEPSC frequency. Acivicin, which decreases efflux of glutamate from the glutathione cycle, decreased mEPSC frequency, a reflection of presynaptic drive. Pyroglutamate restores glutamate, and presynaptic drive (see also Supplementary Figure 2). (*F*) Cumulative probability plots of mEPSC amplitude. Acivicin decreased mEPSC amplitude as well. (#, p<0.01, ‡, p<0.0001 by Steel-Dwass All Pairs tes)t.

We previously reported that in addition to its support of antioxidant function, the glutathione cycle also serves as a reservoir of intracellular neural glutamate (16). Decreasing the liberation of glutamate from the glutathione cycle leads to decreased cortical neuron glutamate, while decreasing the utilization of glutamate increases total glutamate by about 25%. These shifts in glutamate pools could be achieved without increasing oxidative stress or cell death. In the present study we sought to expand this concept by determining if shunting glutamate from the glutathione cycle can shape excitatory neurotransmission. We employed selective inhibitors of different steps of the glutathione cycle and the glutamine-glutamate shuttle and show that glutathione serves as a source for a material portion of glutamatergic neurotransmission.

## Results

### Inhibition of glutathione metabolism depletes neuronal glutamate and affects excitatory transmission

To test the hypothesis that glutathione is a significant reservoir for glutamate, we treated neuronal cells with molecular inhibitors targeting enzymes of the glutathione metabolic cycle: L-2-imidazolidone-4-carboxylate (2I4C), acivicin, L-buthionine sulfoximine (BSO), and sulforaphane. Glutathione and glutamate were quantified by Ellman’s procedure and glutamate oxidase methods (17, 18). If glutathione constitutes a glutamate reservoir, then inhibiting GGT with acivicin should lead to decreased cellular glutamate, as this enzyme is upstream of the ultimate liberation of glutamate from the cycle (Fig. 1*A*, Fig. S1).

As we had previously demonstrated in cell lines (16), total glutamate and glutathione levels declined in primary cortical neurons treated with acivicin (Fig. S2*A-B*). To confirm the specificity of this effect, shRNA targeting of GGT also decreased glutamate levels (Fig. S2*C*). Additionally, we find that the decrease in glutamate brought by acivicin could be rescued by administration of pyroglutamate (5-oxoproline), a downstream metabolite in the glutathione pathway that is a precursor of glutamate (Fig. S1). Pyroglutamate selectively repleted glutamate, but not glutathione (Fig. S2*A-B*).

To determine if glutamate availability from the glutathione cycle (Fig. 1*A*) could shape excitatory transmission, we measured the frequency of miniature excitatory postsynaptic currents (mEPSCs), a reflection of presynaptic drive. Acivicin treatment (24 h) significantly decreased mEPSC frequency (Fig. 1*B, C, E*) and amplitudes to a smaller degree (Fig. 1*B, D, F*). To demonstrate specificity of the effect, we sought to rescue the decreased mEPSC frequency by pretreating with pyroglutamate (PGA), the precursor of glutamate that is synthesized downstream of GGT, which is blocked by acivicin (Fig. 1*A*). Pyroglutamate rescued the effect on mEPSC frequency (Fig. 1*C, E*) but not amplitude, consistent with it being a presynaptic precursor of glutamate. As we previously demonstrated, acivicin at these concentrations did not elicit oxidative stress or affect cell viability (16).

### Inhibition of Glutamate cysteine ligase depletes neuronal glutathione, elevates glutamate and increases excitatory neurotransmission

Glutamate cysteine ligase (GCL) is the rate limiting step for glutathione synthesis, and utilizes glutamate as a substrate. Inhibition of GCL with buthionine sulfoximine (BSO) depleted glutathione rapidly, reflecting its short half-life (1-4 h) as a substrate of multiple enzymes (Fig. S3*B*). BSO treatment increased neuronal glutamate (Fig. S3*A*), consistent with our prior findings in cell lines (16). This was confirmed by shRNA to GCLC, the target of BSO, which increased glutamate levels (Fig. S3*C*). To further test the role of GCL in modulating glutathione and glutamate, we utilized sulforaphane, which increases GCL expression through activation of the Nuclear factor-erythroid 2 p45-related factor 2 (Nrf2) pathway (19). While BSO inhibition of GCL decreased glutathione and increased glutamate, sulforaphane induced GCL, and decreased glutamate while increasing glutathione (Fig. S3*D-F*). We previously demonstrated that these doses and durations of treatment do not alter cortical neuron viability or increase oxidative stress (16). BSO was utilized to test whether increasing the liberation of glutamate from the glutathione cycle could shape excitatory transmission. The BSO-induced increased in glutamate shape excitatory transmission. The BSO induced increased in glutamate was accompanied by an increase in mEPSC frequency (Fig. 2*B, C, E*) and amplitude (Fig. 2*B, D, F*).

**Fig. 2.**
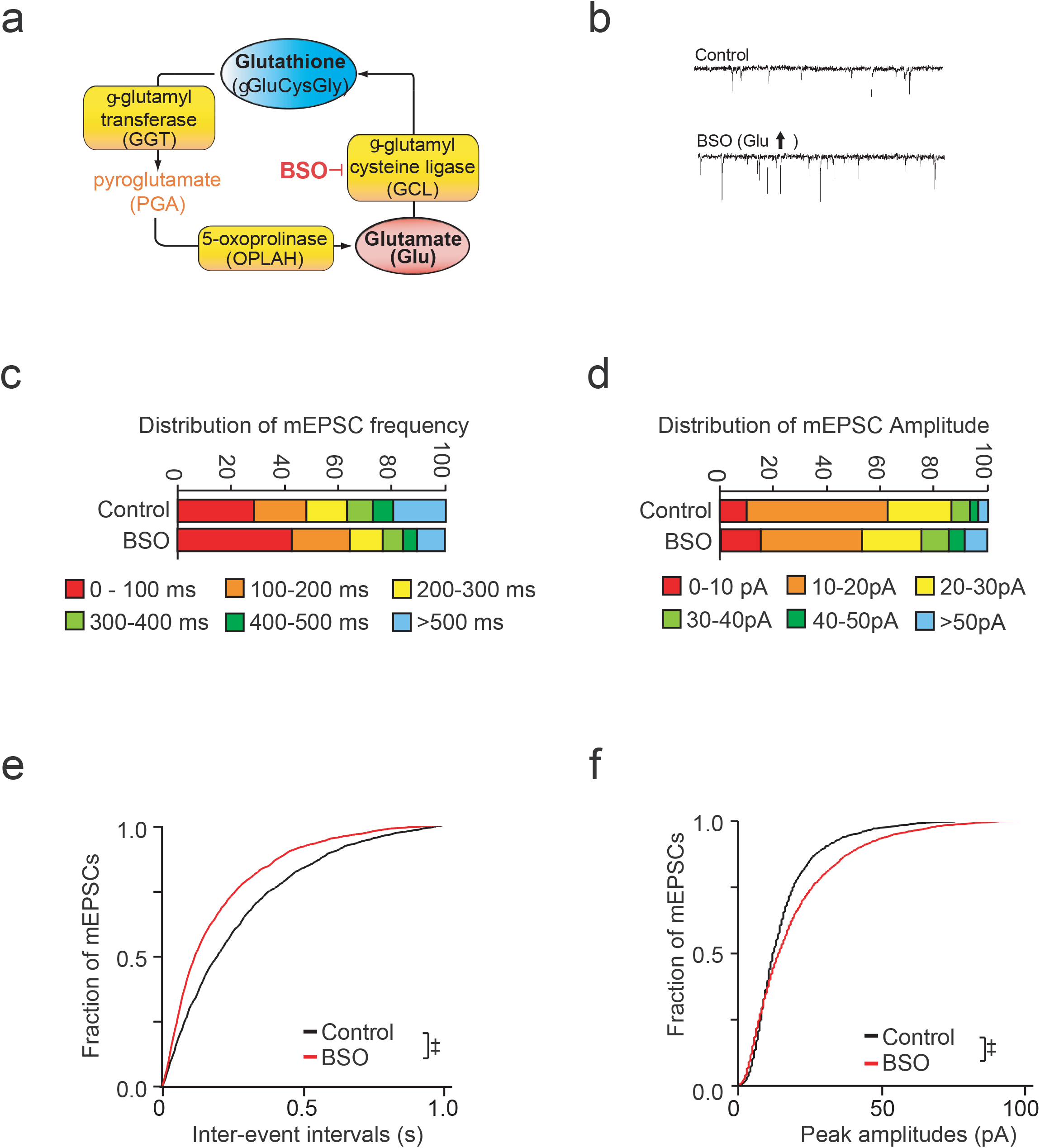
Efflux of glutamate from the glutathione cycle can increase mEPSC. (*A*) Schematic representation of the glutathione cycle and inhibition of glutamate cysteine ligase (GCL) by buthionine sulfoximine (BSO), which shuttles free glutamate into glutathione. Details appear in Supplementary Figures 1 and 4. (*B*) Representative mEPSC traces in primary cortical neurons treated 24 h with vehicle or 200 μM BSO. (*C*) Distribution of mEPSC frequency in cortical neurons treated with BSO. (*D*) Distribution of mEPSC amplitude in cortical neurons treated with BSO. (*E*) Cumulative probability plots of mEPSC frequency. BSO which increases efflux of glutamate from the glutathione cycle, increased mEPSC frequency, a reflection of presynaptic drive (see also Supplementary Figure 4). (*F*) Cumulative probability plots of mEPSC amplitude in cortical neurons treated with BSO. (‡, p<0.0001 by Steel-Dwass All Pairs test).

### The glutathione cycle can complement the glutamate-glutamine shuttle and influence excitatory neurotransmission under conditions of glutamine restriction

The glutamate-glutamine shuttle (Fig. S4) between neurons and glia contributes 50-60% of glutamate neurotransmitter (12, 13, 20) with intracellular sources such as glycolysis supplying the remainder. In the shuttle, glutamate is converted to glutamine in astrocytes, and then exported to neuronal system A transporters, where it is converted to glutamate intracellularly by phosphate activated glutaminase (21). However, glutaminase knockout mice (22) or blockade of system A transporters with methylaminoisobutyric acid (MeAIB) (15) fail to block excitatory neurotransmission. We explored whether the glutathione cycle can complement the actions of the glutamate-glutamine shuttle by blocking import of glutamine by the system A transporter (15), which imports glutamine into neurons, after which it is converted to glutamate (Fig. 3*A*).

**Fig. 3.**
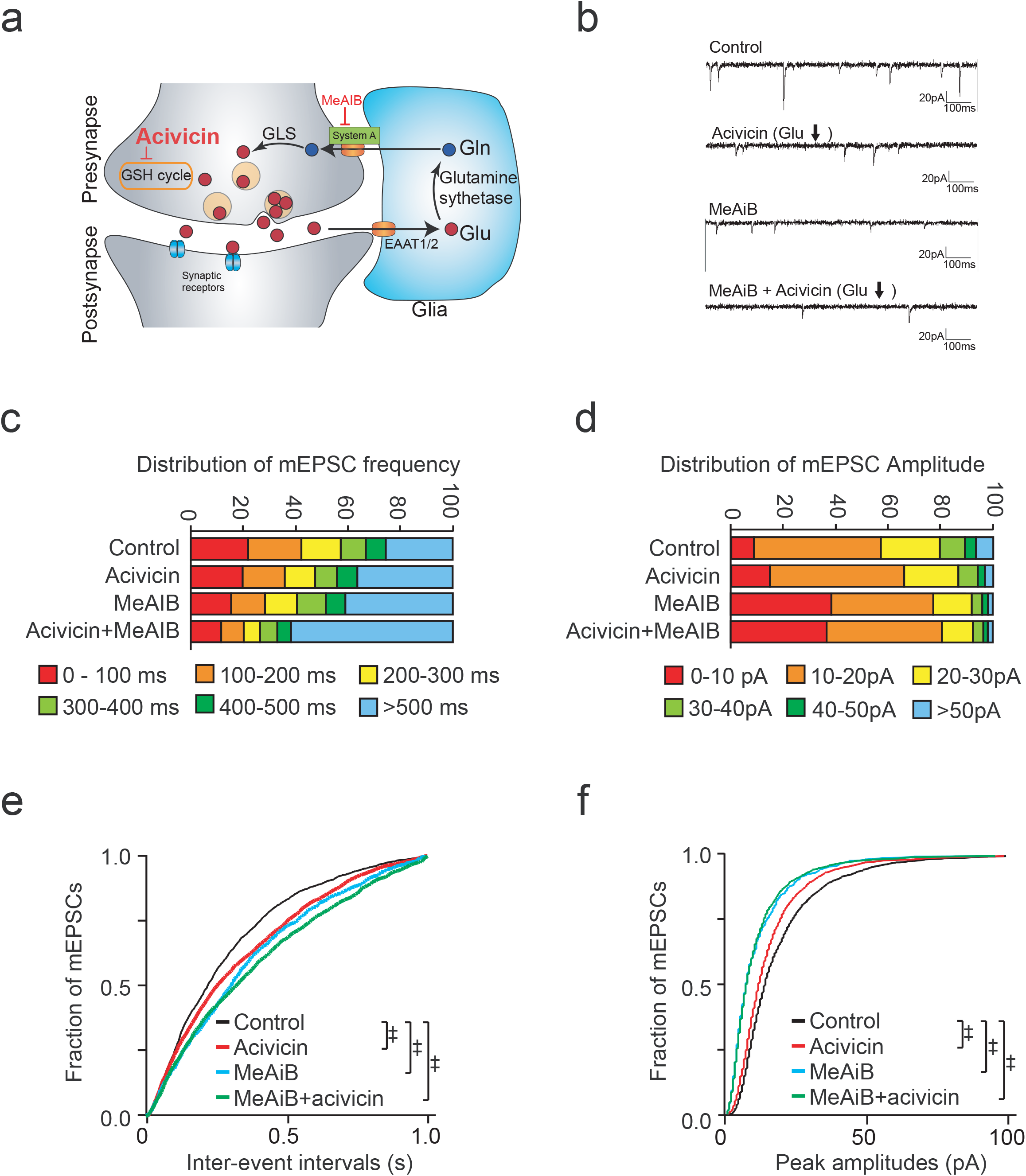
The glutathione cycle supports excitatory transmission, but to a smaller degree than glutamine derived glutamate. (*A*) Scheme of glutamate-glutamine cycling and blockade of system A glutamine transporters by MeAiB. Details appear in Supplementary Fig 6. (*B*) Representative traces of mEPSC recordings in primary cortical neurons treated with 25 μM acivicin (24 h) and/or 25 mM MeAiB (2 h). (*C, D*) Distribution of mEPSC frequency and amplitude in cortical neurons treated with acivicin and/or MeAiB.. (*E, F*) Cumulative probability plots of mEPSC frequency and amplitude. MeAiB decreased mEPSC frequency to a greater degree than acivicin, with both treatments having additive effects for presynaptic drive frequency. (‡, p<0.0001 by Steel-Dwass All Pairs test).

As expected, MeAiB decreased the average mEPSC frequency (Fig. 3*B, C, E*). Acivicin, which diminishes the availability of glutathione-derived glutamate, decreased average mEPSC frequency significantly, though not to the same extent as MeAIB (Fig. 3*E*). mEPSC frequency declined even further when acivicin and MeAIB were co-administered, consistent with glutamate derived from the glutathione cycle having capacity to shape excitatory contributing to maintenance of excitatory activity neurotransmission when glutamine supply is restricted (Fig. 3*C, E*). mEPSC amplitude distributions were similarly diminished by acivicin, and more so by MeAIB, although combinations of acivicin and MeAIB did not further impair the effect of MeAIB alone (Fig. 3*D, F*).

### Glutamate derived from the glutathione cycle rescues excitatory postsynaptic currents during glutamine limitation

We next examined whether glutamate from the glutathione cycle could rescue mEPSC when glutamine supply was restricted (Fig. 4*A*). BSO, which augments glutamate levels, also increased mEPSC frequency (Fig. 2*B, C, E*), while glutamine restriction by MeAIB decreased mEPSC frequency (Fig. 3*B, C, E*). However, administration of BSO could rescue the decreased mEPSC frequency brought by MeAIB (Fig. 4*E*, green line), consistent with the glutathione cycle being induced by MeAiB were rescued by co-administration of BSO, which increased mEPSC frequency (Fig. 4*C,E*), consistent with the glutathione cycle being able to compensate for decreased availability of glutamine, an established source of glutamate neurotransmitter. BSO also significantly improved the decrease in mEPSC amplitudes by MeAIB, though to a smaller degree than its effect upon mEPSC frequency (Fig. 4*D, F*).

**Fig. 4.**
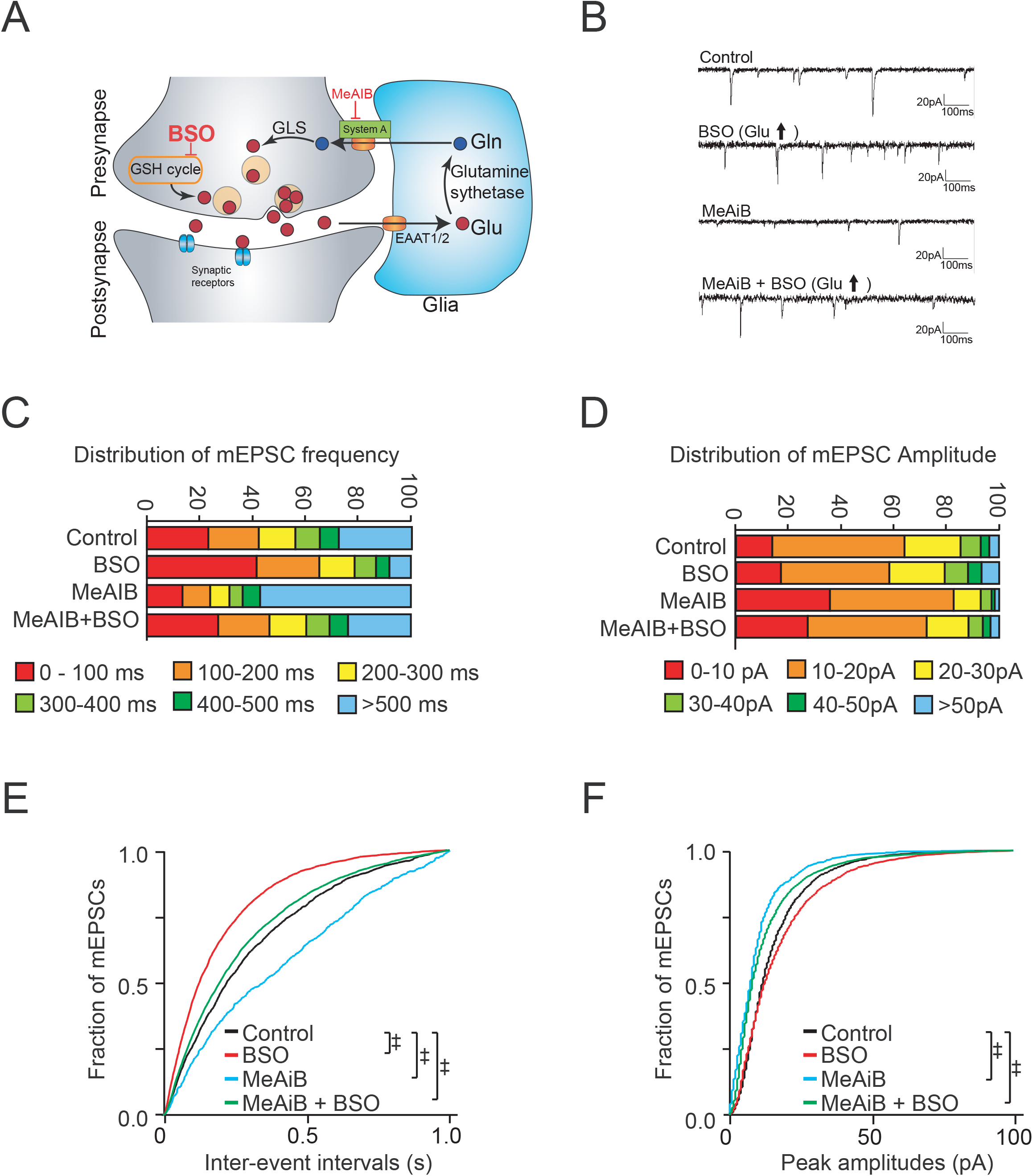
The glutathione cycle can rescue restrictions of glutamine derived glutamate. (*A*) Scheme of glutamate-glutamine cycling and blockade of system A glutamine transporters by MeAiB. Details appear in Supplementary Fig 6. (*B*) Representative traces of mEPSC recordings in primary cortical neurons treated with 25 μM BSO (24 h) and/or 25 mM MeAiB (2 h). (*C, D*) Distribution of mEPSC frequency a nd amplitude in cortical neurons treated with BSO and/or MeAiB. (*E, F*) Cumulative probability plots of mEPSC frequency and amplitude. MeAiB, which restricts glutamine derived glutamate, inhibits mEPSC frequency (blue). Decreases in mEPSC frequency induced by were rescued by pre-treatment with BSO (green), which increased mEPSC frequency (red). BSO also improved the decrease in mEPSC amplitudes by MeAIB. (‡, p<0.0001 by Steel-Dwass All Pairs test).

## Discussion

We previously reported that the glutathione cycle may serve as a reservoir of total neuronal glutamate (16). Treatments that decrease the liberation of glutamate from the glutathione cycle lead to decreased intracellular glutamate, whereas decreasing utilization of glutamate or glutathione synthesis increases it (16). We now report that shunting of glutamate derived from glutathione can shape excitatory neurotransmission. Inhibiting GCL, which utilizes glutamate to synthesize glutathione, leads to increased glutamate and mEPSC frequency. Decreasing the release of glutamate from the glutathione cycle by blocking GGT leads to diminished mEPSC. We have previously demonstrated that these fluxes of cytosolic glutamate can occur without detectable increases in oxidative stress or altered cell viability (16). Thus the glutathione pathway is poised to contribute glutamate without impacting neuronal viability unless there are massive, sustained deficits (16, 23). Our study was designed to test the effect of modulating glutathione cycle metabolism and its effect on glutamatergic activity. The effects on EPSC frequency were associated with respective increases or decreases in intracellular glutamate. We demonstrated specificity of the effect of acivicin, which decreases glutamate liberation from glutathione, by rescuing with pyrogluamate, the immediate metabolic precursor of glutamate in the glutathione cycle. Another mode by which GSH could potentially modulate neurotransmission is by its effects on redox-regulated presynaptic proteins such as synaptosomal-associated protein 25 (SNAP-25) and N-ethylmaleimide-sensitive factor (NSF). These changes could be operational in neurodegenerative diseases, reflecting an imbalance in redox balance modulated by GSH. The redox effects of GSH or other thiol agents can be assessed by the use of irreversible thiol alkylating agents such as N-ethyl maleimide (NEM) which would form covalent bonds with cysteine residues and prevent further modification of these sites as has been analyzed in modulation of GABA- and glycine-evoked currents in rat retinal ganglion cells (24).

The glutathione pathway may also supply glutamate when glia derived glutamine is blocked by MeAIB, which inhibits system A transporters. While glia-derived glutamine provides 50-60% of glutamate neurotransmitter, when the pathway is blocked with MeAIB excitatory transmission abruptly decreases but rapidly recovers, consistent with endogenous neuronal sources of glutamate neurotransmitter (15). We suggest glutathione is one such endogenous source, as further impairing glutamate liberation from the cycle diminishes synaptic activity induced by MeAIB, while increasing glutamate availability can rescues the impairment by MeAIB. A reservoir capacity of glutathione may be utile during periods of sustained synaptic activity.

Alterations in neuronal glutamate levels, such as by fluxes to and from a glutathione reservoir, might have an impact upon glutamate neurotransmitter. Vesicular glutamate transporters have a much lower affinity for glutamate 0.5-3.5 mM than plasma membrane transporters GLT1/EAAT2, whose Km is 4-40 μM (25, 26). Furthermore, as glutamate neurotransmitter typically does not saturate postsynaptic receptors, modest impacts upon release frequency may influence synaptic strength (27-30). Our findings also affirm prior reports that increasing intracellular glutamate concentration in presynaptic terminals leads to greater excitatory post synaptic currents (EPSC) (29, 30). In our specific approach, we suggest that the glutathione cycle may be one such source of this glutamate. This may provide some mechanistic implications to interpret magnetic resonance spectroscopy techniques in human subjects, in which total regional brain glutamate may be determined (31, 32). 7 Tesla (7T) proton magnetic resonance spectroscopy (MRS) studies have shown that glutamate levels were significantly lower in first episode psychosis (FEP) subjects, whereas glutamine levels were unaltered (33). This study also revealed lower levels of glutathione in the anterior cingulate cortex (ACC) and thalamus, which supports the idea of origin of glutamate from glutathione (16). Metabolic flux analysis utilizing ^13^C labeled glutathione would yield additional information regarding flux of glutathione and glutamate under various conditions. Use of "Phasic” axons that fatigue in their glutamate neurotransmitter release have lower glutamate levels than “tonic” glutamate axons with greater glutamate levels (34). Glutamine levels are similar in both, suggesting that significant reservoirs of glutamate exist in neurons independent of glutamine. Localized glutathione synthesis would be expected to have an even more pronounced effect, and it has been suggested that non-soma areas contain more glutathione (35).

These findings may be relevant to human disease. Glutamatergic dysfunction has been implicated in schizophrenia by multiple lines of evidence (36-43). Several investigators have reported aberrant glutathione levels in schizophrenia patients, including medication naïve subjects (44-48). Mice lacking the modifier subunit of GCL, have a 60% reduction in glutathione, accompanied by abnormal cortical gamma synchrony, decreased parvalbumin interneurons (PV-IN) and behavioral phenotypes relevant to schizophrenia.(49-51). Despite substantial glutathione deficits, the mice are outwardly healthy. Additionally, rare deficiencies of glutathione cycle enzymes (gamma-glutamylcysteine ligase, glutathione synthetase, 5-oxoprolinase, and gamma-glutamyl transferase) have all been associated with neuropsychiatric and cognitive impairments, although detailed phenomenological characterization has not been reported. (9, 52, 53). A role for glutathione as a glutamate reservoir may be a bridge between distinct lines of research that implicate glutamatergic dysfunction and aberrant glutathione levels in neuropsychiatric conditions such as schizophrenia.

We suggest that two drugs available for human use, sulforaphane, which increases glutathione, and pyroglutamate, which is converted to glutamate in the glutathione cycle, may be therapeutically beneficial. Sulforaphane (54) is a potent inducer of the Nrf2 transcription factor, has excellent blood brain barrier penetration (55), and might expand the size of the glutathione reservoir by increasing expression of GCL, the rate liming step in glutathione biogenesis. Recent studies in human subjects show that sulforaphane elevates glutathione levels and those of other brain metabolites (56). Sulforaphane has also been reported to improve symptoms of autistic spectrum disorder (57). Pyroglutamate is a glutamate precursor whose CSF concentration is 120 μM (58), rivaling the extracellular glutamine concentration of 400 μM (basal glutamate is 2-3 μM). Oral administration of pyroglutamate has been found to benefit age-associated memory impairment (59), alcoholic encephalopathy (60), and delirium induced by anticholinergic medication (61). Pyroglutamate may be a promising therapeutic candidate for cognitive dysfunction in schizophrenia and other conditions with glutathione disturbances.

## Acknowledgements

The authors are indebted to Dr. Minori Koga for experimental support and discussions. The authors wish to sincerely thank Yukiko Lema for assistance. Support by NIH NINDS 1K08NS057824 and Johns Hopkins BSI and NARSAD Young Investigator Award to TWS., National Institute of Mental Health (MH-092443 Silvio O. Conte center, MH-094268, NIH MH 084018, MH-105660, and MH-107730, as well as foundation supports from the Stanley and S/R to A.S.

## Author contributions

T.W.S., B.D.P., A.S, S.H.S. designed the research. G.M.P., B.D.P., and Y.T conducted experiments. S.H.S., A.S and T.W.S. directed the research. T.W.S., B.D.P., S.H.S. and A.S analyzed data. B.D.P, S.H.S., A.S and T.W.S. wrote the paper.

## Materials and Methods

### Cell culture and reagents

Dissociated cortical neuron cultures from Sprague-Dawley rats were prepared as described previously (62). Primary cortical neurons were maintained in Neurobasal medium^®^ (Life Technologies Corporation) supplemented with 1x B-27 (life technology). Cells were maintained at 37 °C in 5% CO2/95% atmosphere and medium replaced every 3 days. For glutathione and glutamate quantification, cell viability analysis and oxidative stress assay, cells were sub-cultured in 12-well plates at a density of 4.2 × 10^5^ cells/well. For electrophysiological recording, cells were seeded onto 12 mm diameter poly-D-lysine coated glass coverslips at a density of in 24-well plates and grown for 48 hours before experiments. The media was then replaced with fresh media and treated with 25 μM acivicin, 100 mM 2-imidazolidone-4-carboxylate (2I4C), 200 μM buthionine sulfoximine (BSO) or 5 μM pyroglutamate (PGA) for glutathione and glutamate measurement. Alpha-methylaminoisobutyric acid (MeAIB) was treated 1 to 4 hours before the electrophysiological recording. Acivicin was obtained from BIOMOL. 2I4C, BSO and PGA were obtained from Sigma. MeAIB was purchased from Chem-Impex International. All animal procedures related to were approved by the Johns Hopkins University Animal Care and Use Committee.

### Measurement of glutathione

Total and oxidized glutathione was determined by the Tietze method with minor modifications (63). Cells were washed in PBS, scraped and suspended in phosphate buffer then sonicated. A portion of lysate was suspended in 0.1% N-lauroylsarcosine and used for analysis of protein content by Bradford assay (64). The remaining solution was centrifuged 15 minutes at 20,000g and the soluble fraction was used for detection of glutathione content. Proteins in the remaining fraction were precipitated with 50 mg/mL metaphosphoric acid, removed by centrifugation, and supernatants neutralized with 200 mM triethanolamine. For oxidized glutathione measurement, an aliquot of the supernatant was incubated 60 minutes at room temperature with 10 mM 2-vinylpyridine (2-VP) to scavenge reduced glutathione. The rate of increase in absorbance at 415 nm, which measures the reduction of 5-5’-dithiobis (2-nitrobenzoic acid) by glutathione, reflects the total glutathione content (or oxidized glutathione when 2-VP is added). The concentration of total and oxidized glutathione content in cells was calculated by a calibration curve with standards.

### Measurement of glutamate

Cells were collected in phosphate buffer (100 mM sodium phosphate, 1 mM ethylene diamine tetraacetate, pH 7.5) and sonicated and lysates centrifuged to remove insoluble debris. Determination of glutamate was performed by Kusakabe’s method (65). 2-10 nmol of L-glutamate standards and the cell lysate were mixed with reaction buffer (final concentrations of 36.8 mM 4-(2-hydroxyethyl)-1-piperazineethanesulfonic acid (HEPES) buffer (pH 7.1), 2.19×10-5 U L-glutamate oxidase, 1 U peroxidase, 0.8 mM 3.5-dimethoxy-N-ethyl-N-(2-hydroxy-3-sulphopropyl) aniline (sodium salt) (DAOS), 0.8 mM aminoantipyrine). After incubating at 22 °C for 30 minutes, the mixture was measured at a wavelength of 570 nm with a microplate spectrophotometer. The concentration of L-glutamate was then calculated using a standard calibration curve, normalized to protein sample concentrations, and levels expressed as percentage relative to controls.

### mEPSC recordings

Whole-cell patch clamp recordings were performed at 22 °C from primary cortical neurons at days *in vitro* of 12-16. Coverslips containing the neurons were loaded into an upright microscope (Olympus BX-61-WI) fitted with a Warner RC-26 submerged recording chamber. Patch pipettes (4-6 Mohm) were filled with cesium solution containing 115 mM cesium methanesulfonate, 20 mM 4-(2-hydroxyethyl)-1-piperazineethanesulfonic acid, 10 mM disodium phosphocreatine, 5 mM tetraethylammonium chloride, 3 mM adenosine 5’-triphosphate magnesium, 2.8 mM sodium chloride, 0.5 mM guanosine 5’-triphosphate sodium, and 0.4 mM ethylene glycol tetraacetic acid (pH 7.2, 285-290 mOsm/kg). Recordings were made at room temperature in artificial cerebrospinal fluid (aCSF) (pH 7.3, osmolarity 305 +/- 5 mOsm/kg) perfused at 2 mL/min containing 150 mM NaCl, 10 mM 4-(2-hydroxyethyl)-1-piperazineethanesulfonic acid, 10 mM glucose, 3 mM potassium chloride, 2 mM calcium chloride, and 1.3 mM magnesium chloride. Putative neurons were identified under visual guidance with infrared differential interference contrast optics (Olympus BX-61-WI) with a 40x water-immersion objective. The image was captured with an infrared-sensitive EMCCD camera (IXON DU885K) and displayed on a monitor. In addition, the image was sent to a computer with a serial connector and captured (Metamorph Advanced). Whole-cell current clamp recordings were made with a headstage (CV-7B) connected to a computer-controlled amplifier (MultiClamp 700B, Axon Instruments), digitized through a Digidata 1440A Analog/digital converter and acquired at a sampling rate of 10 kHz. All current steps to set membrane potential and elicit action potentials were delivered through the recording pipette and controlled by Clampex 10 (Axon). Electrode potentials were adjusted to zero before recording. mEPSCs were recorded in the presence of tetrodotoxin (1 μM, Tocris, Ellisville, MO, USA or Alomone, Jerusalem, Israel) and picrotoxin (0.1mM, Sigma-Aldrich, St. Louis, MO, USA). All voltage clamp recordings were performed at a holding potential of −70mV. mEPSC events were detected using the software MiniAnalysis (Synaptosoft, Decatur, GA, USA) with a 5 pA amplitude threshold and all mEPSCs were verified visually. Data for analysis was excluded if access resistance is more than 50 Mohm or membrane resistance is less than 100 Mohm. Cumulative probability histograms were tested by Kolmogorov-Smirnov Test for each recording from at least 50 events. Mean values for mEPSC frequency and amplitude between groups were analyzed by Tukey-Kramer HSD test. All results are shown as mean +/- S.E.M.

#### Isolation of synaptosomes

Synaptosomes were isolated as described previously (66). Briefly, mice were euthanized by decapitation and whole brains homogenized in 20 vol. of 0.32 M sucrose. After centrifugation at 1000 x g for 10 min, the supernatant (S1) was centrifuged for 20 min at 17.500 x g to yield the supernatant S2 and the pellet P2 (crude synaptosomal fraction). P2 was suspended in 10 vol of 0.32 M sucrose and 10 ml of the solution layered on a 0.8 and 1.2 M two-step discontinuous gradient, which was centrifuged at 61,000 x g for 2 h to separate the synaptosomes from the mitochondria (which forms the pellet). The synaptosomes form the interface between the 1.2 M and 0.8 M sucrose layers.

### Statistical analysis

All data were expressed as the mean+/-S.E. Statistical differences in glutathione, glutamate measurement and miniature EPSC analysis were done by Tukey-Kramer HSD test, except where noted. Statistical differences between two groups for behavioral tests were determined with Student’s t test, and statistical differences among three groups or more were determined using a one-way analysis of variance (ANOVA), two-way ANOVA, and an ANOVA with repeated measures, followed by Bonferroni’s multiple comparison test; p<0.05 was regarded as statistically significant.

## Supporting Information

**Fig. S1.**
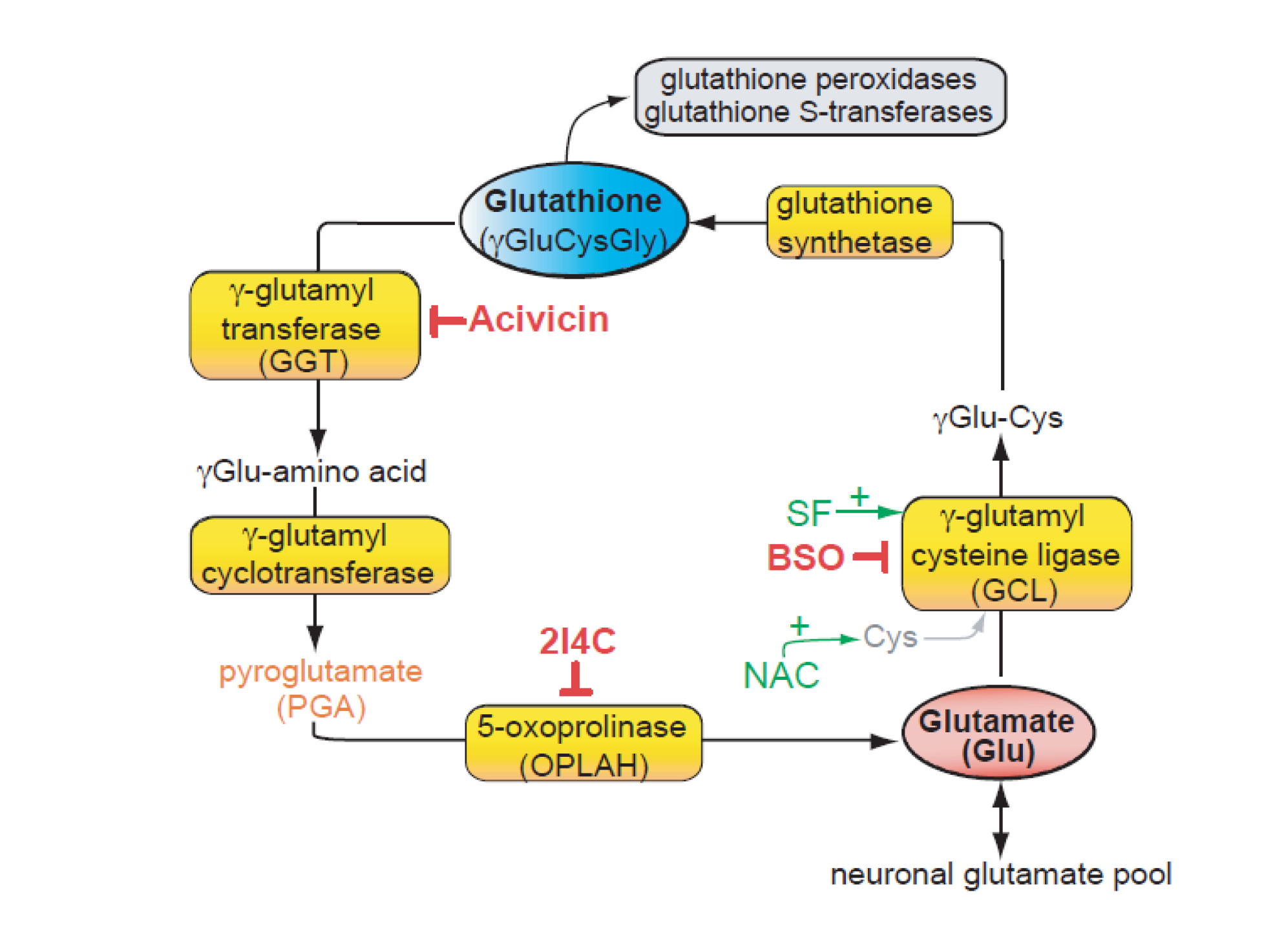
The gamma-glutamyl cycle (glutathione cycle). Glutamate is used as a substrate for glutathione synthesis by GCL and glutathione synthetase. Glutamate is liberated from glutathione through the action of GGT, glutamyl cyclotransferase and OPLAH. Inhibitors (acivicin, 2I4C, and BSO) are shown with their respective targets. PGA is a metabolite of GGT and precursor of glutamate in this cycle. OPLAH, 5-oxoprolinase; GCL, gamma-glutamyl cysteine ligase; 2I4C, 2-imidazolidone-4-carboxylate; BSO, buthionine sulfoximine; NAC, N-Acetylcysteine; SF, sulforaphane; PGA, pyroglutamate.

**Fig. S2.**
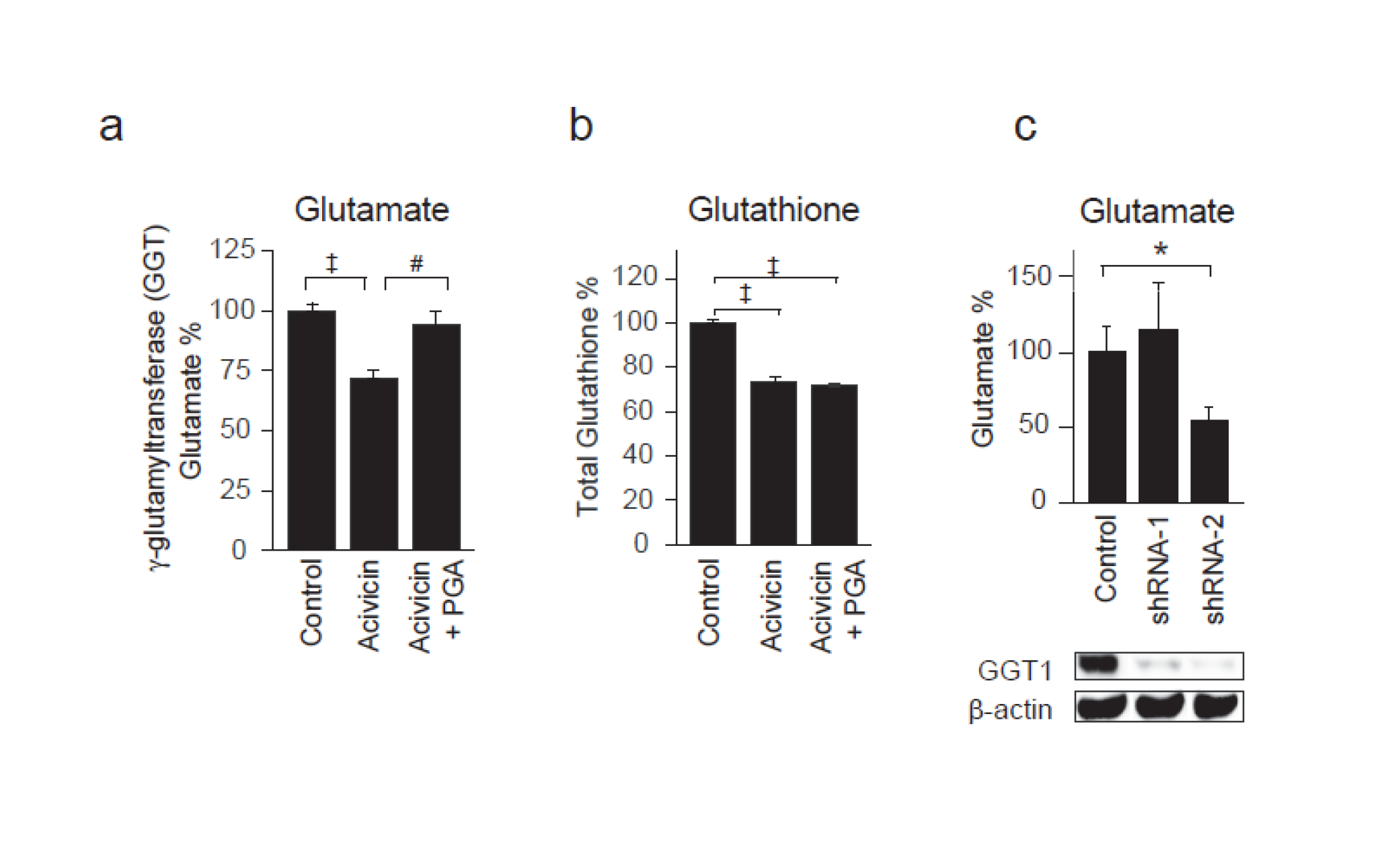
Inhibition of γ-glutamyltransferase diminishes glutamate levels. (*A,B*) Acivicin treatment of cortical neurons (25 μM, 24 h) decreases glutamate and glutathione levels. Co-administration of 5 μM pyroglutamate (PGA) selectively rescues the glutamate decrease. (*C*) shRNA targeting of γ-glutamyltransferase in N2A neurons decreases glutamate. shRNA-2 had greater suppression of protein expression than shRNA-1. *, p<0.05, #, p<0.01, † p<0.001, ‡ p<0.0001 by 1-way ANOVA with Tukey-Kramer posthoc.

**Fig. S3.**
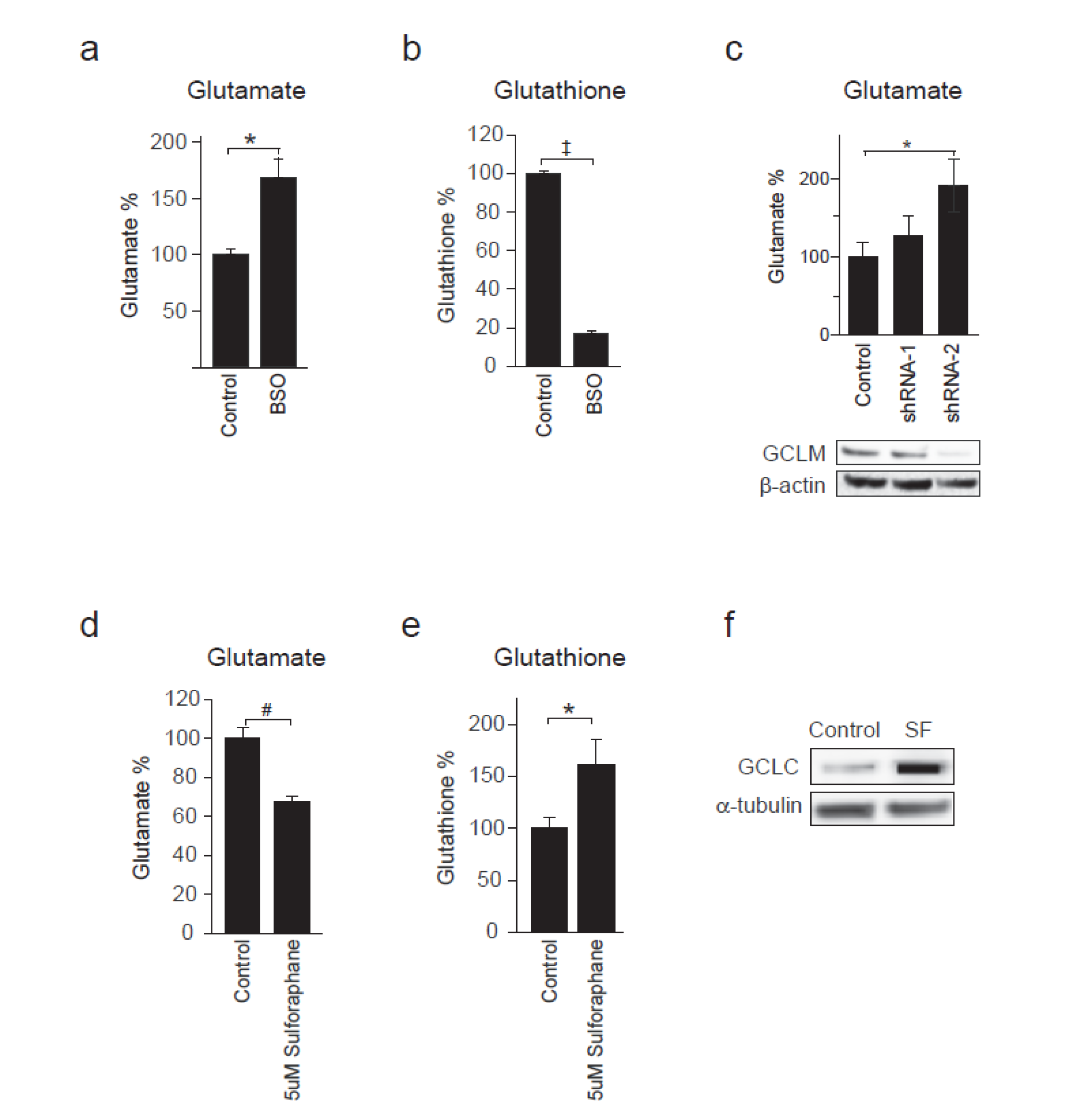
Targeting GCL to modulate glutathione and glutamate levels. (*A,B*) BSO, which blocks GCL, decreases glutathione and increases precursor glutamate. Cortical neurons were treated 24 h with 200 μM BSO (*A,B*), or N2A neurons were treated with shRNA to GCL (*C*). shRNA-2 gave greater suppression of GCL protein than shRNA-1. (*D-F*) Sulforaphane increases GCL and acutely increases glutathione and decreases glutamate. Cortical neurons were treated 24 h with 5 μM sulforaphane. Lysates were immunoblotted with anti-GCLC antibody. BSO, buthionine sulfoximine; GCL, glutamate-cysteine ligase. *, p<0.05, #, p<0.01, † p<0.001, ‡ p<0.0001 by 1-way ANOVA with Tukey-Kramer posthoc.

**Fig. S4.**
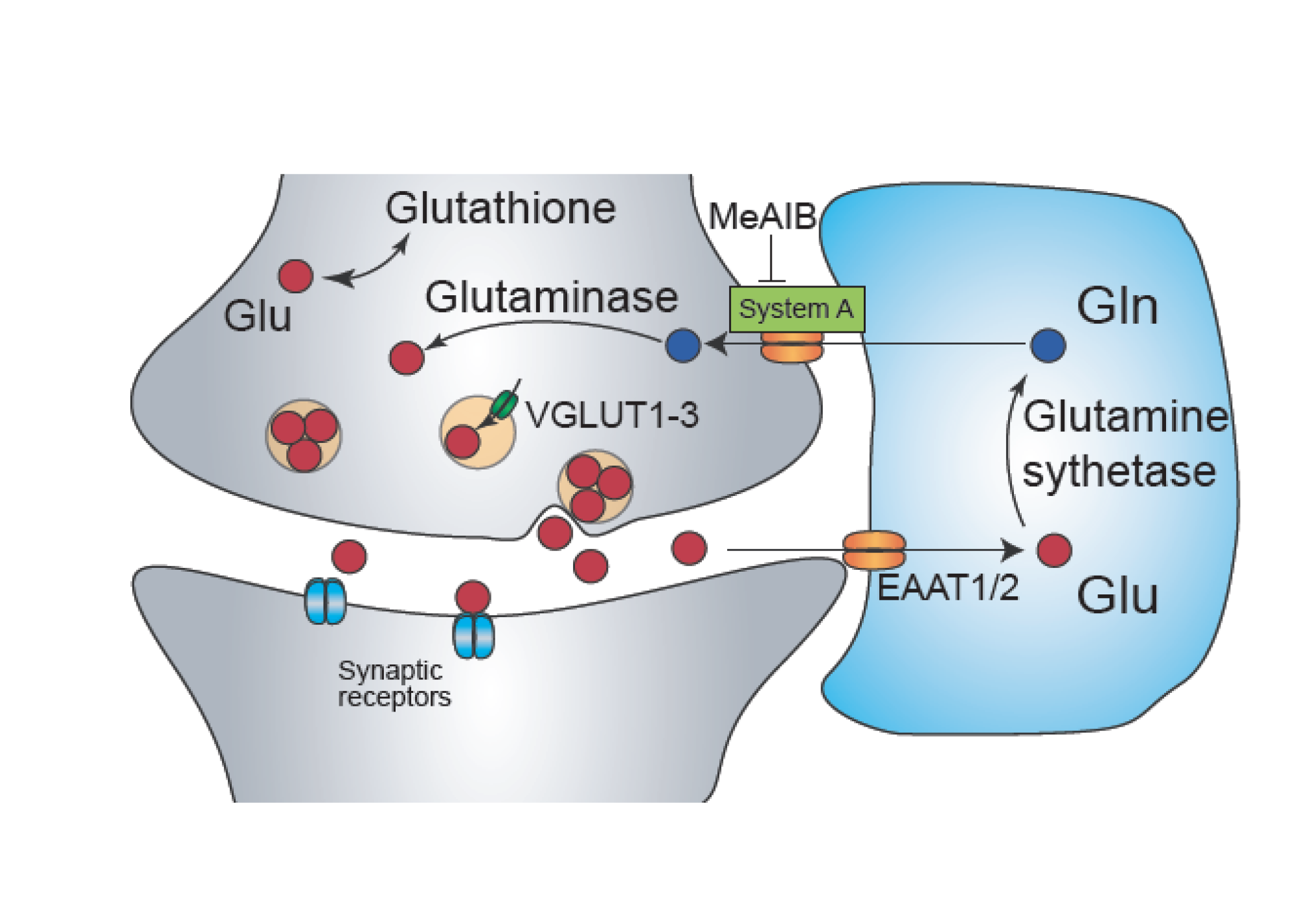
Model for the glutathione cycle complementing the glutamine-glutamate shuttle. Schema of cycling of glutamate-glutamine and glutamate-glutathione. Solid red circles indicate glutamate and solid blue circles indicate glutamine. MeAiB, blocks system A glutamine transporters that import glutamine.

**Fig. S5.**
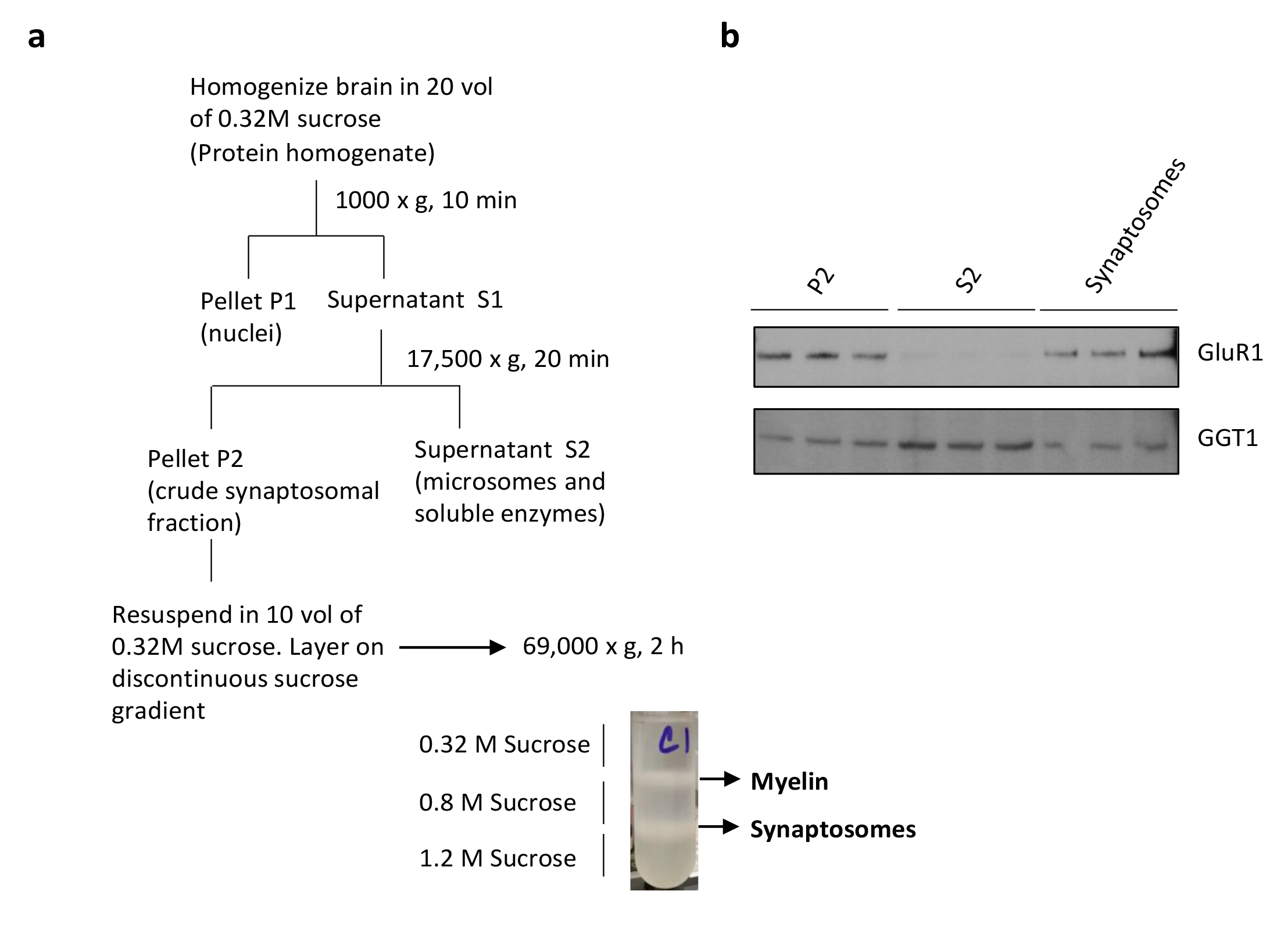
Gamma-glutamyl transferase (GGT) is associated with the synaptosomal fraction. (*A*) Subfractionation protocol to isolate synaptosomes. After discontinuous sucrose density gradient centrifugation, the synaptosomes band at the interface of the 0.8 M and 1.2 M sucrose layers. (*B*) Western blot showing localization of GGT1 to the synaptosomal fraction (n=3). GGT1 is also present in the soluble fraction, S2. GluR1 (the AMPA receptor) was used as a positive control. GluR1 is present only in the synaptosomal fractions and absent from the soluble fraction, S2.

